# Temporal response characterization across individual multiomics profiles of prediabetic and diabetic subjects

**DOI:** 10.1101/2021.12.08.471816

**Authors:** Minzhang Zheng, Carlo Piermarocchi, George I. Mias

**Affiliations:** Michigan State University, Biochemistry and Molecular Biology, East Lansing, MI 48824, USA; Michigan State University, Institute for Quantitative Health Science and Engineering, East Lansing, MI 48824, USA; Michigan State University, Physics and Astronomy, East Lansing, MI 48824, USA

## Abstract

Longitudinal deep multi-omics profiling, which combines biomolecular, physiological, environmental and clinical measures data, shows great promise for precision health. However, integrating and understanding the complexity of such data remains a big challenge. Here we propose a bottom-up framework starting from assessing single individuals’ multi-omics time series, and using individual responses to assess multi-individual grouping based directly on similarity of their longitudinal deep multi-omics profiles. We applied our method to individual profiles from a study profiling longitudinal responses in type 2 diabetes mellitus. After generating periodograms for individual subject omics signals, we constructed within-person omics networks and analyzed personal-level immune changes. The results showed that our method identified both individual-level responses to immune perturbation, and the clusters of individuals that have similar behaviors in immune response and which was associated to measures of their diabetic status.

## Introduction

The development of novel technologies in personal health monitoring devices, high throughput sequencing and computational methods has generated massive omics data, and provides both a great opportunity and challenge to precision health [1–5]. The big data provides plentiful health information ranging from biomolecular, physiological, and environment data to clinical measures. This information helps identify potential deviations from a healthy baseline and improve health risk predictions [1]. A big challenge of a big data approach to precision health is how to integrate and understand these multi-dimensional, extremely diverse sources, with highly heterogeneous data [3]. Early efforts by Chen, Mias, Li-Pook-Than et al. focused on assessing the feasibility integrated Personal Omics Profiling (iPOP), by utilizing a multiomics integration framework to interpret healthy and diseased states followed through an individual’s blood-based multiomics assessment [6]. More recent efforts by Sara Ahadi et al. revealed personal aging markers by using deep longitudinal profiling [7], Abdellah Tebani et al. discovered how the personal cohort changes during the wellness period [8]. Environmental effecs have also been studied by M. Reza Sailani et al., revealing two biological seasonal patterns in California by multi-omics profiling [9]. Wearable sensors are also be used in digitalized health in tracking physiomes and activity [10]. Other implementations have used multi-omics to monitor the drug responses [11]. Non-invasive longitudinal saliva multi-omics has been recently used by Mias et al. to monitor immune responses in a vaccinated individual [12]. Although these efforts have shown the great promise of deep multi-omics profiling, the complexity of data presents limitation for practical implementations. Deep multi-omics data come from diverse sources, and have different types, sizes and ranges, which complicates comparisons between different individuals’ personal multi-omics. In the Pioneer study study by Price et al [13] dynamic data clouds were used for longitudinal monitoring of individual subjects, in a study that also incorporated behavioral coaching to improve clinical biomarkers. In their recent work [14] carried out iPOP across multiple individuals, and built correlation networks of molecular associations. However, to the best of our knowledge, direct networks of individuals associated with longitudinal individual deep multi-omics profiles have not been constructed. In this work, we present a framework to categorize the personal longitudinal deep multi-omics profile and group the individuals into communities, using spectral representations of individual multi-omics time series. We provide a new approach to compare the longitudinal deep multi-omics profile between different individuals regardless of the data heterogeneity. We applied our method to personal multi-omics profiling data from prediabetic or diabetic individuals (Type 2 diabetes mellitus, T2D) at its earliest stage from the study by Zhou et al [14]. We identified individual-level responses to immune perturbation, and clusters of individuals showing similar behavior. The microscopic molecular behavior was linked to phenotypic differences, including body mass index and insulin resistance, with the immune response dominating differences attributed to diabetic status.

## Results

### Summary of cohort details and data

The original data used in this analysis comes from the study by Zhou et al. [14], that focused on multi-omics characterization of host-microbe dynamics in prediabetics. The different subjects in the study had highly heterogeneous visits records: some subjects only had one visit record but one subject has more than 150 visit records (time points), which is about 15 times more than the average of 10 visits per subject. Multiple omics were generated from the subjects including blood based transcriptomics, microbiome data (nares/gut), cytokine measurements and clinical measures. To ensure the individual omics profiles had enough time points and time series, we filtered records from individuals so that the number of time points *N_t_* >= 4 and the number of omics *N_o_* > 500, across participants from the data source [14]. We also excluded the subject with the 150+ visits records, as this was not comparable directly to other subjects in our analysis, given the density of points. Our final filtered dataset contained 69 subjects from the original data source. Figure 1A shows the cumulative bar graph of age and sex distribution of the subjects. We had 35 males and 34 females, and most subjects were older than 50. A histogram of the subject’s observation window is shown in Figure 1B. The observation window is heterogeneous, ranging from 200 to 1200 days. During the observation window, there were total 846 visits with different healthy conditions, including: 486 healthy visits as baseline, 148 visits when subjects got infected, 119 visits had immunization effects, 43 visits with subject weight gain or loss period, 18 visits with subjects on antibiotics, and 32 other healthy conditions, as summarized in Figure 1 C. Overall we analyzed 733,425 time series across 5 datasets: 713,874 RNA-sequencing (RNA-seq) data, 3,221 clinical measures, 4,554 cytokines, 6,336 Gut and 5,440 Nares measurements. The majority (>90%) of the time series comes from RNA-seq, Figure 1D.

**Fig 1.**
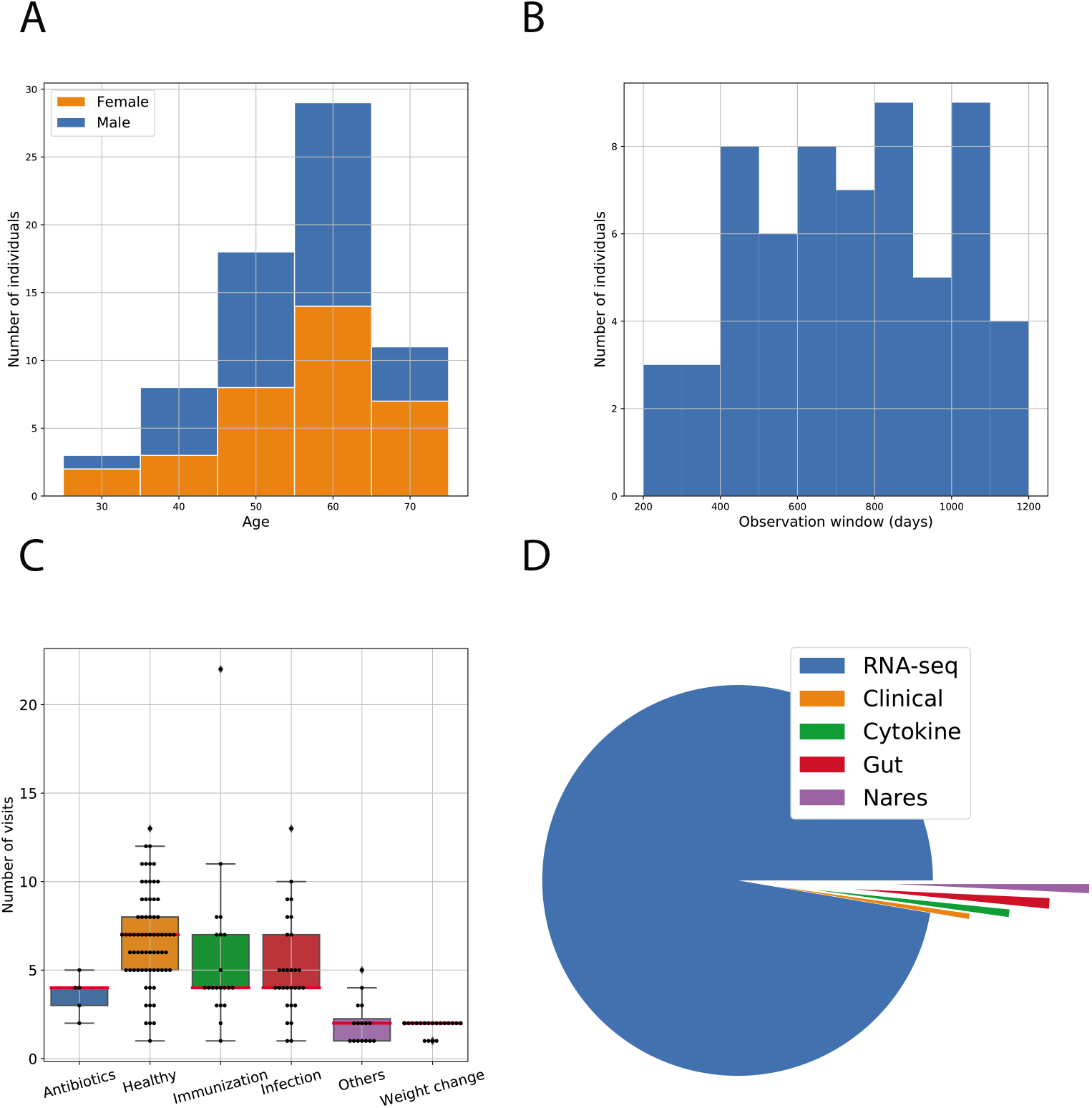
Cohort description. (A) The cumulative bar graph of subjects’ sex and age distribution. (B) Histogram of observation window. (C) Distributions of visits and conditions. (D) Proportion of time series from different data modalities.

### Single subject analysis

We used PyIOmica’s [15] spectral methods to classify the time series for each individual’s omics into temporal trends. The objective is to identify sets of omics that show similar temporal behavior that deviates from each individual’s own baseline. The PyIOmica categorization algorithm generates 3 sets of classes from an individual’s omics time series: (i) *Lag classes*, of time series showing statistically significant autocorrelation at different lags, (ii) *Spike Maxima class* of time series with no autocorrelation but with positive spikes (high intensity pulses), and (iii) *Spike Minima class* of time series with no autocorrelation but with negative spikes (low intensity pulses) [15,16]. Then within each class, the algorithm separates time series into groups and subgroups based on the autocorrelation patterns and signal intensities. Further details are provided in the Methods.

All analysis and results from each individual’s classifications are provided in the Online Data Files (ODFs). Here we show examples of Lag1 classification for two subjects, ZKFV71L (Female, 66-year old, Prediabetic) and ZTMFN3O (Female, 40-year old, Prediabetic), Figure 2(A) and (B) respectively. The Lag 1 class for subject ZKFV71L has 323 time series, which were assigned to 1 group and 3 subgroups, shown in Figure 2(A) left panel. The intensity changes of the 3 subgroups represented the healthy status changes, indicating the systemic immune responses in the subject. Within each subgroup, we created a mean time series, whose intensities of each time point equals to the average intensity of the time series at this time point within the subgroups. These are used to obtain the time points community structure using our visibility-based community detection algorithm [17]. The communities of each subgroup were related to the health status changes, as indicated in Figure 2 middle panel (circular visibility graph layouts are shown for each subgroup’s mean time series). The autocorrelation heatmap corresponding to the time series in this category is also shown in Figure 2 right panel. Similar results were found in subject ZTMFN30, Figure 2(B). Additional details, including corresponding omics in each class and group/subgroup classification are available in the ODFs for all subjects.

**Fig 2.**
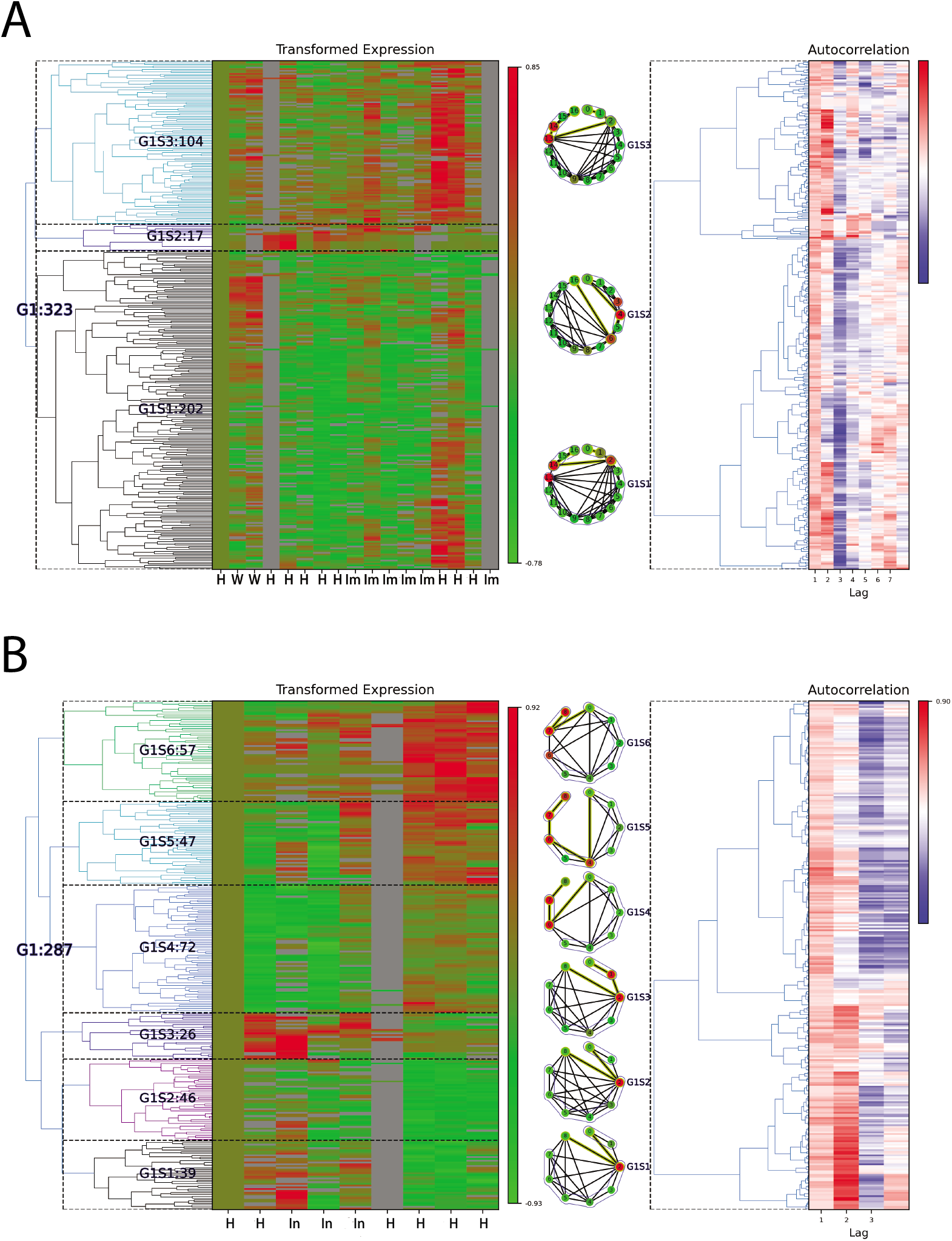
Single individuals’ multi-omics clusters. Two examples of Lag 1 classification outcomes are shown for (A) Subject ZKFV71L. (B) Subject ZTMFN3O. In these examples the information is summarized as follows: *Left panel*: the cluster of groups/subgroups for Lag 1 class are shown in the visits time frame. The visit time points have been labeled by healthy status, where H: Healthy, W: Weight gain/loss, Im: Immunization, In: Infection. *Middle panel*: the community structure of visits within each subgroup, where the community structure is based on our visibility-graph-based community detection algorithm [17]. *Right panel*: corresponding autocorrelations for the time series shown.

Once sets of omics that show similar profiles in time are identified, we can assess the biological significance of these temporal associations. Following classification, we carried out Reactome pathway [18] enrichment analysis for the genes that showed statistically significant trends for each subject. The statistically significant (False Discovery Rate, *FDR* < 0.05) pathways results for subject ZKFV71L and ZTMFN3O for the autocorrelation Lag 1 class are shown in Table 1. The over-representation for subject ZKFV71L included Endosomal/Vacuolar pathway (14 genes), Antigen Presentation: Folding, assembly and peptide loading of class I MHC (15 genes), Interferon alpha/beta signaling (16 genes), ER-Phagosome pathway (14 genes), Interferon gamma signaling (16 genes), Antigen processing-Cross presentation (14 genes), Immunoregulatory interactions between a Lymphoid and a non-Lymphoid cell (15 genes), Interferon Signaling (16 genes), Class I MHC mediated antigen processing & presentation (17 genes) and Adaptive Immune System (23 genes), etc. These Reactome pathways indicated immune responses of this subject (ZKFV71L) corresponding to the health status change from Healthy to Immunization. Similarly, we also found statistically significant Reactome pathways for subject ZTMFN3O, including: Cellular response to heat stress (7 genes), Cellular responses to stress (12 genes), Cellular responses to stimuli (12 genes), Interleukin-10 signaling (7 genes), Signaling by Interleukins (13 genes) and Cytokine Signaling in Immune system (14 genes). These pathways are also indicative of an immune response of this subject (ZTMFN3O) corresponding in this case to a health status change from Healthy to Infection. Additional Reactome Pathway enrichment analyses for all subgroups and for all subjects are included in the ODFs.

**Table 1.** Statistically significant (*FDR* < 0.05) Reactome pathways results for subject ZKFV71L and ZTMFN3O for autocorrelation Lag 1.

| Reactome pathway | Matched IDs | p value | FDR |
| --- | --- | --- | --- |
| **A. Subject ZKFV71L** |  |  |  |
| Endosomal/Vacuolar pathway | 14 | 1.10E-13 | 3.05E-11 |
| Antigen Presentation: Folding, assembly and peptide loading of class I MHC | 15 | 1.17E-13 | 3.05E-11 |
| Interferon alpha/beta signaling | 16 | 4.90E-11 | 8.48E-09 |
| ER-Phagosome pathway | 14 | 1.82E-09 | 2.37E-07 |
| Interferon gamma signaling | 16 | 3.32E-09 | 3.45E-07 |
| Antigen processing-Cross presentation | 14 | 8.14E-09 | 7.00E-07 |
| Immunoregulatory interactions between a Lymphoid and a non-Lymphoid cell | 15 | 4.82E-07 | 3.56E-05 |
| Interferon Signaling | 16 | 1.49E-06 | 9.71E-05 |
| Class I MHC mediated antigen processing & presentation | 17 | 3.52E-06 | 0.000200 |
| Adaptive Immune System | 23 | 9.36E-05 | 0.00487 |
| RUNX3 regulates RUNX1-mediated transcription | 2 | 0.000718 | 0.0337 |
| **B. Subject ZTMFN3O** |  |  |  |
| Attenuation phase | 7 | 2.415E-10 | 5.121E-08 |
| HSF1-dependent transactivation | 7 | 1.152E-09 | 1.221E-07 |
| Regulation of HSF1-mediated heat shock response | 7 | 9.528E-08 | 6.670E-06 |
| Cellular response to heat stress | 7 | 3.135E-07 | 1.462E-05 |
| HSF1 activation | 5 | 3.482E-07 | 1.462E-05 |
| Cellular responses to stress | 12 | 4.469E-05 | 0.002 |
| Cellular responses to stimuli | 12 | 5.365E-05 | 0.002 |
| RMTs methylate histone arginines | 3 | 7.407E-04 | 0.019 |
| Interleukin-10 signaling | 7 | 2.416E-07 | 8.408E-05 |
| Signaling by Interleukins | 13 | 1.283E-05 | 0.0022 |
| NR1H3 & NR1H2 regulate gene expression linked to cholesterol transport and efflux | 4 | 0.00032 | 0.0353 |
| Signaling by Nuclear Receptors | 8 | 0.00057 | 0.0353 |
| Cytokine Signaling in Immune system | 14 | 0.00073 | 0.0353 |
| Signaling by Overexpressed Wild-Type EGFR in Cancer | 2 | 0.00073 | 0.0353 |
| Inhibition of Signaling by Overexpressed EGFR | 2 | 0.00073 | 0.0353 |
| NR1H2 and NR1H3-mediated signaling | 4 | 0.00083 | 0.0353 |
| NR1H2 & NR1H3 regulate gene expression to limit cholesterol uptake | 2 | 0.00114 | 0.0353 |
| NR1H2 & NR1H3 regulate gene expression linked to gluconeogenesis | 2 | 0.00114 | 0.0353 |
| NR1H2 & NR1H3 regulate gene expression linked to triglyceride lipolysis in adipose | 2 | 0.00114 | 0.0353 |
| EGFR interacts with phospholipase C-gamma | 2 | 0.00137 | 0.0357 |
| Interleukin-18 signaling | 2 | 0.00137 | 0.0357 |

### Multi-subject similarity analysis

Based on the individual results, we first aggregated the omics that showed statistically significant trends in each individual to identify the signals that are consistent across the majority of individuals (>50% of subjects, number of occurrences ≥35), included in Table 2. We found that high frequency signals came from 3 data sources: cytokines, nares and gut microbiome. Many studies have now shown that cytokines have a profound relationship with Type 2 diabetes [19–21], our findings are consistent with these previous works and provide potential biomakers for Type 2 diabetes (see also Discussion).

**Table 2.** Frequency of signals with statistically significant temporal trends in individuals.

| **ID** | **Source** | **# Occurrences** |
| --- | --- | --- |
| genus_Streptococcus | nares | 43 |
| IP10 | cytokine | 43 |
| class_Gammaproteobacteria | nares | 42 |
| family_Streptococcaceae | nares | 39 |
| phylum_Proteobacteria | gut | 38 |
| IL13 | cytokine | 38 |
| PDGFBB | cytokine | 37 |
| class_Bacilli | gut | 35 |
| family_Streptococcaceae | gut | 35 |
| genus_Streptococcus | gut | 35 |
| order_Lactobacillales | gut | 34 |
| class_Betaproteobacteria | nares | 34 |
| genus_Dorea | gut | 33 |
| TGFA | cytokine | 33 |
| IL27 | cytokine | 33 |
| TGFB | cytokine | 33 |
| phylum_Bacteroidetes | nares | 33 |
| genus_Blautia | gut | 33 |
| ALKP | clinical | 32 |
| family_Micrococcaceae | nares | 32 |
| MCP1 | cytokine | 32 |
| genus_Flavonifractor | gut | 32 |
| class_Bacteroidia | nares | 32 |
| order_Bacteroidales | nares | 32 |
| family_Propionibacteriaceae | nares | 32 |
| genus_Propionibacterium | nares | 32 |
| genus_Clostridium.XIVa | gut | 32 |
| GCSF | cytokine | 31 |
| MIG | cytokine | 31 |
| genus_Oscillibacter | gut | 31 |
| IL22 | cytokine | 31 |
| genus_Clostridium.IV | gut | 31 |
| CD40L | cytokine | 30 |
| VEGF | cytokine | 30 |
| order_Lactobacillales | nares | 30 |
| SCF | cytokine | 30 |
| EGF | cytokine | 30 |
| family_Coriobacteriaceae | gut | 30 |
| order_Coriobacteriales | gut | 30 |

To further investigate common responses across individuals, we created a multi-subject similarity network. The network was constructed by comparing the Euclidean distance of the spectral time series representation (periodogram) across common omics between every pairs of subjects. The network, shown in Figure 3(A), has nodes corresponding to the 69 subjects, with weighted edges corresponding to the number of omics that showed similar temporal behavior. We used a k-means algorithm to calculate communities of the network (see Methods section for details). Three communities were identified, denoted as Community 0, Community 1, and Community 2. Community 0 has 44 individuals, 24 Females and 20 Males, ages from 29 to 75, with disease status including 33 Prediabetic, 1 Diabetic, 7 Crossover and 3 Control. Community 1 has 15 individuals, 6 Females and 9 Males, ages 33 to 71, 11 Prediabetic, 1 Crossover and 3 Control. Community 2 has 10 individuals, 5 Females and 5 Males, age from 39 to 70, 6 Prediabetic, 2 Crossover and 2 Control. We then compared clinical measures between the subjects in the community, including BMI, disposition index (DI), steady-state plasma glucose (SSPG), Matsuda index (Matsuda), and maximum insulin secretion rate (isrMax). The violin plots show group separated by community and sex, Figure 3(B)-(F). These 5 distribution figures qualitatively indicate that the Community 0 and Community 1 have similar distribution but have large differences with community 3. We also found that the female and male subjects have different distributions, even within same community for BMI, DI and isrMax measures. We carried out non-parametric Mann-Whitney U tests [22,23] to compare across the different communities for statistical significance (p-value < 0.05), with the results summarized in Table 3. Most of the 5 measures do not show statistical significant differences between Community 0 and Community 1, except for isrMax comparison between males in Community 0 and Community 1. The BMI and SSPG distributions between Community 0 and Community 2, and between Community 1 and Community 2 have statistically significant differences, indicating that the subjects in Community 2 have statistical difference in physiological states comparing with Community 0 and Community 1. Differences in Male vs. Females in the comparisons, particularly for “BMI”, “DI” and “isrMax”, suggest that even though overall subjects in these two communities may have similar physiological states and responses, females and males still display different physiological states (though the low number of subjects is affecting power in further breaking down of community differences).

**Fig 3.**
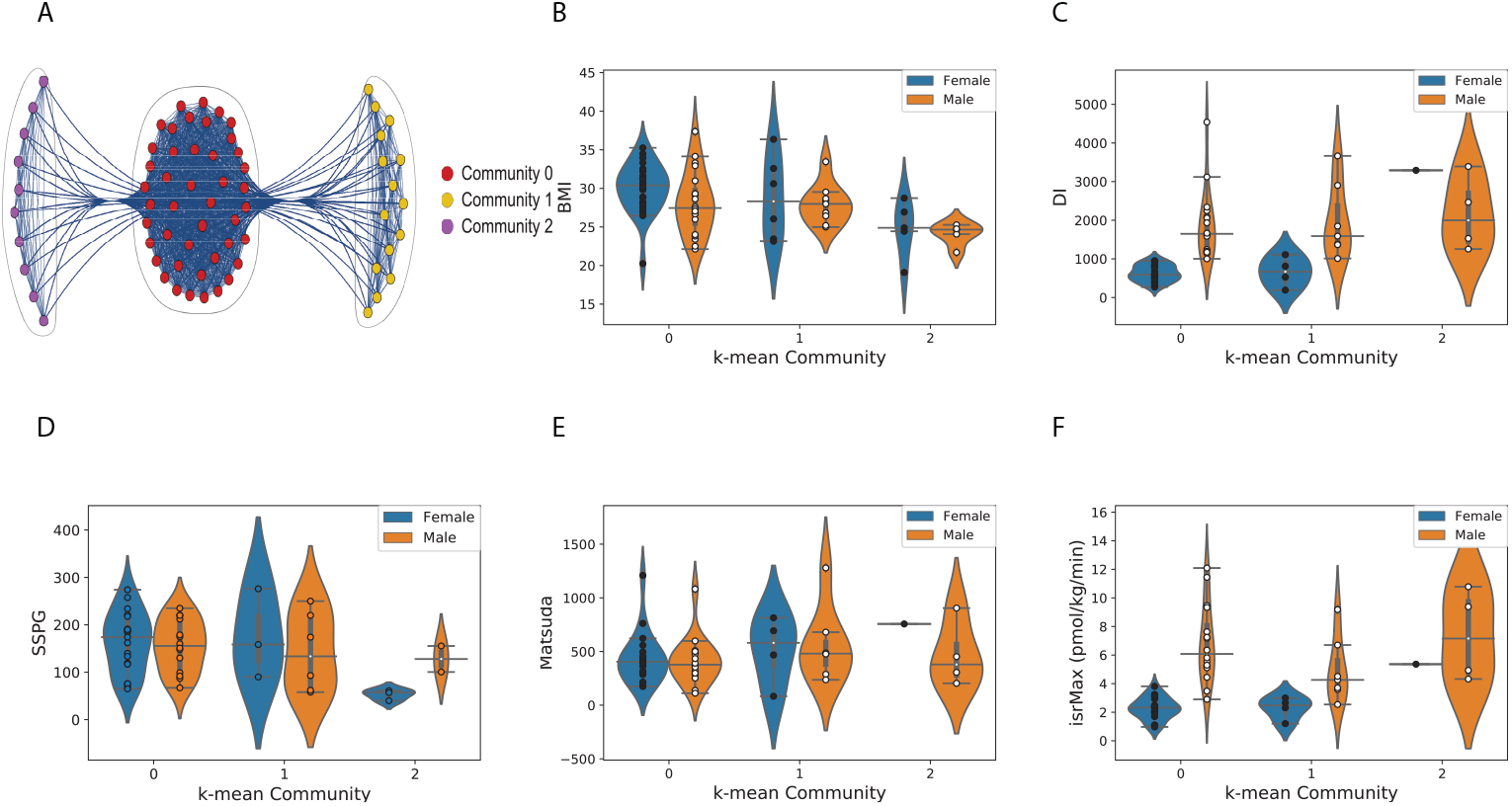
Similarity analysis across individuals. (A) the k-means based community structure of the subjects’ similarity network (nodes represent subjects and weighted edges omics showing similar temporal behavior across individuals). (B) - (F) Distributions of 5 types of measures in the 3 network communities by gender: (B) BMI; (C) Di, disposition index; (D) SSPG, steady-state plasma glucose; (E) Matsuda index and (F) isrMax, maximum insulin secretion rate. Community 0 and Community 1 have similar distributions, but these two communities show differences in distributions compared to Community 2.

**Table 3.** Mann–Whitney U test for different measures between two different communities. The labels C0, C1, C2 correspond to Community 0, Community 1 and Community 2, respectively. Statistically significant (p-value < 0.05) results are shown in red.

| Comparison | BMI | SSPG | Matsuda Index | Disposition Index | isrMax |
| --- | --- | --- | --- | --- | --- |
| C0 vs C1 Female | 0.28 | 0.42 | 0.20 | 0.45 | 0.42 |
| C0 vs C1 Male | 0.49 | 0.36 | 0.18 | 0.46 | 0.044 |
| C0 vs C1 Total | 0.14 | 0.36 | 0.086 | 0.16 | 0.45 |
| C0 vs C2 Female | 0.0030 | 0.0037 | 0.11 | 0.060 | 0.060 |
| C0 vs C2 Male | 0.022 | 0.35 | 0.48 | 0.21 | 0.48 |
| C0 vs C2 Total | 0.00034 | 0.0054 | 0.27 | 0.0050 | 0.033 |
| C1 vs C2 Female | 0.21 | 0.040 | 0.36 | 0.14 | 0.14 |
| C1 vs C2 Male | 0.0038 | 0.43 | 0.25 | 0.46 | 0.054 |
| C1 vs C2 Total | 0.0080 | 0.042 | 0.45 | 0.070 | 0.012 |

We next ranked the omics in each community (represented by the weighted edges), by their frequencies of occurrence. We then carried out Reactome Pathway Enrichment analysis for the top 50% ranked genes of each community (to reduce noise effects from low frequency genes). The statistically significant pathway results (FDR < 0.05) are shown in Table 4. Community 0 has 11 statistical significant Reactome pathways, related to immune response. For Community 1 the statistically significant Reactome pathways include Translocation of ZAP-70 to Immunological synapse. Community 2 did not have statistically significant Reactome results. The Reactome pathway Enrichment analysis, along with the subjects’ BMI, DI, SSPG, Matsuda and isrMax distributions differences indicate that BMI, SSPG and the isrMax affect temporal immune responses, and also have sex differences. In summary, our methodology can separate individuals with different physiological conditions and immune response based on the multi-omics time series similarity of individual profiling analyses.

**Table 4.** Statistically significant (*FDR* < 0.05) Reactome pathways results of top half highest genes for Communities 0 and 1.

| Reactome pathway | Matched IDs | p value | FDR |
| --- | --- | --- | --- |
| **A. Community 0** |  |  |  |
| Antigen Presentation: Folding, assembly and peptide loading of class I MHC | 35 | 1.110E-16 | 1.077E-14 |
| Endosomal/Vacuolar pathway | 35 | 1.110E-16 | 1.077E-14 |
| Interferon alpha/beta signaling | 38 | 1.110E-16 | 1.077E-14 |
| ER-Phagosome pathway | 36 | 1.110E-16 | 1.077E-14 |
| Antigen processing-Cross presentation | 36 | 1.110E-16 | 1.077E-14 |
| Interferon gamma signaling | 36 | 1.110E-16 | 1.077E-14 |
| Immunoregulatory interactions between a Lymphoid and a non-Lymphoid cell | 36 | 1.110E-16 | 1.077E-14 |
| Interferon Signaling | 39 | 7.772E-16 | 6.528E-14 |
| Class I MHC mediated antigen processing & presentation | 38 | 1.062E-12 | 7.968E-11 |
| Cytokine Signaling in Immune system | 48 | 4.642E-07 | 3.110E-05 |
| Adaptive Immune System | 43 | 3.855E-06 | 2.351E-04 |
| **B. Community 1** |  |  |  |
| PD-1 signaling | 15 | 1.686E-12 | 1.898E-09 |
| Translocation of ZAP-70 to Immunological synapse | 14 | 9.325E-12 | 5.250E-09 |
| Phosphorylation of CD3 and TCR zeta chains | 14 | 2.287E-11 | 8.576E-09 |
| Generation of second messenger molecules | 15 | 7.107E-11 | 1.997E-08 |
| Costimulation by the CD28 family | 15 | 5.149E-08 | 1.158E-05 |
| MHC class II antigen presentation | 17 | 4.332E-07 | 8.100E-05 |
| Downstream TCR signaling | 15 | 1.104E-06 | 1.766E-04 |
| TCR signaling | 16 | 1.875E-06 | 2.625E-04 |

## Discussion

In this manuscript we have applied spectral methods to analyze multiomics individual profiles from public data for 69 individuals. The goals were to detect deviations from an individual’s own baseline for each subject, as well as compare results across subject in a bottom-up approach. We have generated periodograms for individual subject omics time series categorization, and constructed within-person omics networks and analyzed personal-level immune changes. We the used periodograms across individuals to identify network clusters of individuals with similarities across their common omics temporal patterns.

Our analysis discovered both intra- and inter-individual characteristics differences. The inter-individual longitudinal omics clusters show pattern changes corresponding to the individual’s physiological state changes. We identified similar individual-level responses to immune perturbation. The multi-individuals’ similarity network revealed different classes within which the molecular behavior was linked to phenotypic differences, including body mass index and insulin resistance, with the immune response dominating differences attributed to diabetic status, Table 3.

Previous research has reported several cytokines with important roles in Type 2 diabetes development, including cytokines from our findings across individuals in Table 2, with some examples highlighted below. Elevated concentrations of IP10 (CXCL10) have been reported in type 2 diabetes, and associated with higher diabetes risk [24, 25]. The IL13 pathway is a potential therapeutic target for glycemic control in type 2 diabetes [26]. IL27 has been implicated in insulin resistance in genome wide association studies [27]. Wang et al. had reported the pathogenic role of IL27, using diabetic NOD mice to investigate T-cell mediated autoimmune diabetes [28], but recently the role of IL27-IL27R*α* in promoting adipocyte thermogenesis has been investigated in the context of treating insulin resistance [29]. Decreased plasma IL22 level was found to be a potential trigger of impaired fasting glucose and type 2 diabetes, in a retrospective study of Han Chinese subjects [30]. PDGFBB is reported associate with Type 2 diabetes mellitus and complications [31,32]. TGFA [33] and TGFB [34] have shown pathologic contribution in diabetic kidney disease. TGFB is also reported associated with Type 2 diabetic nephropathy [35]. ALKP has been investigated as an independent predictor for diabetes incidence [36,37]. MCP1 has been found significantly increased in patients with type 2 diabetes [38]. Furthermore, through rat studies GCSF has been reported as a potential novel therapeutic drug in early diabetic nephropathy patients [39]. The involvement of MIG (CXCL9) in the progression of type 2 diabetes nephropathy has been reported [40]. CD40-CD40L has been associated with type 2 diabetes mellitus [41], VEGF is involved in the pathogenesis of diabetic complications [42], c-Kit and its ligand, stem cell factor (SCF) have been reported as a potential novel target for treating diabetes [43]. Finally, chemotactic cytokines, including eosinophil chemotactic factors (ECFs), have been shown to be related to T2D [44]. In our results, the cytokines above had high occurrences rates with statistically significant trends across the diabetic and prediabetic individuals. The findings suggest that our approach has potential application in clinical trials to identify disease biomakers and treatment targets.

Our study has limitations. The transcriptomic data used were generated from bulk RNA-sequencing, and hence do not allow for cell-type specific analyses, which are important is evaluating Type 2 diabetes. We expect that cell-type-specific studies with longitudinal data will become more prevalent with the recent focus on single-cell RNA-sequencing approaches. In terms of the data, there is uneven sampling (inherent in any real-world/subject based study), which we have addressed with our approach, as well as having different lengths of time series across individuals. Furthermore, environmental measures are not included and the low number of participants does not allow for a nuanced analysis of heterogeneity across subjects. For example, different subjects received different treatments which are analyzed in bulk. In terms of broad applicability of data collection and analysis, the data are obtained using invasive approaches (at least for blood components), which should improve with non-invasive transcriptomic mapping, such as using saliva. While our analysis included microbiome data (nares and gut), their association with transcriptome results, the cytokines and other measurements noted in Table 2, and their mechanistic role still requires further model-based experimentation.

In this manuscript we analyzed previously obtained data from an analysis on prediabetes [4, 14], which revealed insights from this longitudinal dataset and can lead to actionable health discoveries, providing relevant information for precision health monitoring. Our approach is the first, to our knowledge, to implement a temporal bottom-up methodology. Starting from individual microscopic measurements (molecular level omics), intra-subject immune responses can be characterized. Then, building on the individual responses, a macroscopic inter-subject temporal clustering of subjects based on temporal similarity provides information on how immune responses can be related to diabetic states, consistent with the original work. In summary, our findings utilize personal temporal omics to identify collective responses across individuals associated with macroscopic characteristics, and provide an approach paradigm that can potentially help predict disease responses and outcomes towards clinical implementations.

## Methods

### Data preprocessing

The data used in this manuscript came from personal multi-omics profiling data from individuals with Type 2 diabetes mellitus at its earliest stage [14]. The measures, SSPG, Matsuda, DI and isrMax, came from the other paper of the same project [4].

To obtain an individual’s omics profile, we combined all the omics source dataset into a dataframe, then separated into dataframes for each individual. Since our workflow has two branches: single subject analysis and multi-subject similarity analysis, seen in Figure 4, the following data preprocessing was carried out for the two branches: (i) For the single subject analysis, we selected the signals with less than 25% time points from each individual’s dataframe as the input for single subject analysis, using each individual’s time points as possible measurement points. (ii) For multi-subject analysis, we sorted each individual’s time frame from their dataframes, then combined all individual time frames to get all the possible time points as the common time frame. We then calculated Lomb-Scargle periodograms from each individual’s dataframe using this common time frame as the set of possible measurement points. The transformed dataframe was then used as the input for the multi-subject analysis.

**Fig 4.**
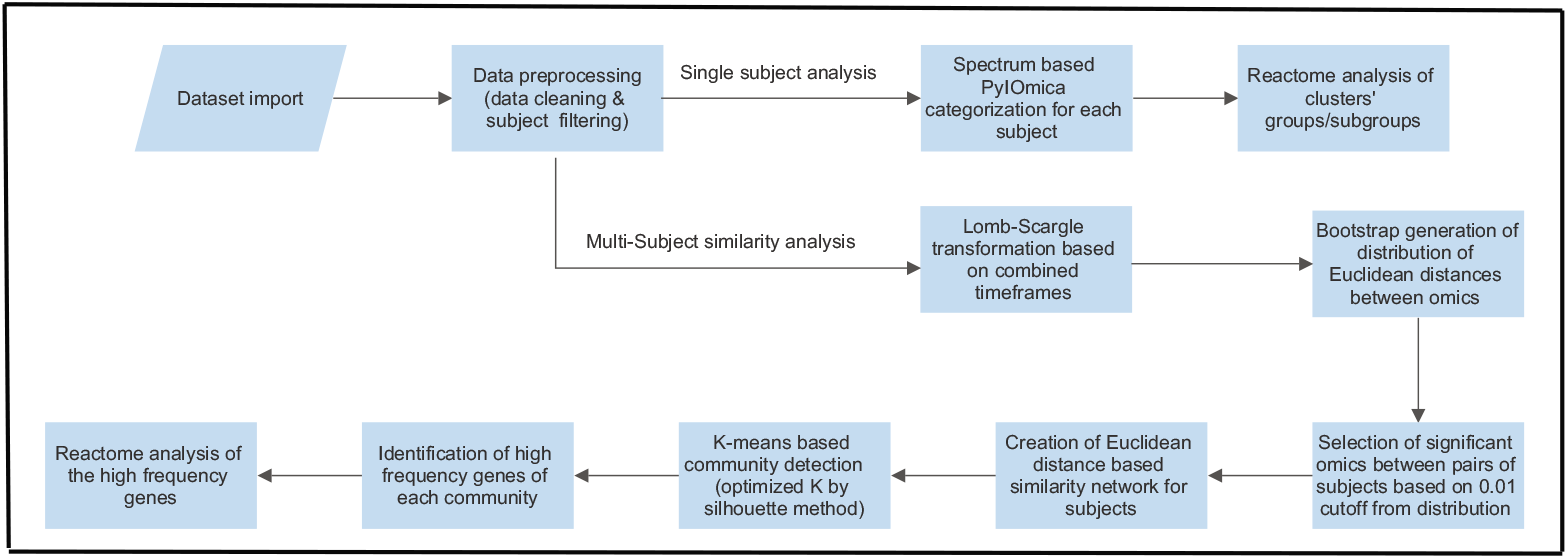
Algorithmic flowchart. Our workflow has two main branches: (i) single subject analysis and (ii) multi-subject similarity analysis.

### Individual subject analysis

#### Time series categorization

The individual subject analysis was carried out in Python using PyIOmica’s `calculateTimeSeriesCategorization` command [15]. Briefly, for each subject *s*, for each omics *i* its time series *X* were analyzed over its constituent timepoints. The individual omics intensities at timepoint *j* were compared to the initial timepoint by subtracting its intensity from all values, 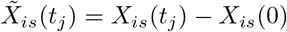. The data were then normalized to a time series *Q*, using the Euclidean norm, so that

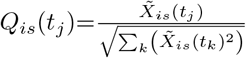

The algorithm’s classification of signals into trends uses spectral methods, as previously described [12, 15]. Briefly, for each signal a Lomb-Scargle periodogram is calculated as a list *P_i_*. The inverse Fourier transform of *P_i_* returns a list of autocorrelations, {*ρ_ik_*}, where *k* ∈ {0,…, *N/2*} is the lag. In parallel with the original time series signals, a bootstrap set of 10^5^ time series were generated by sampling from the original data with replacement. The autocorrelations at different lags of the bootstrap set were computed to generate an autocorrelation null distribution for each lag from which a set of cutoffs {*ρ_ck_*} were obtained corresponding to a 0.95 quantile. A time series was then assigned to a class labeled with the lowest *Lag l* for which the series’ autocorrelation *ρ_il_* is larger than the cutoff, i.e., where *l* = Min [{*j*: *ρ*_ij_ ≥ *ρ*_cj_}], and *j* ∈ 1,…, *k*.

If a time series 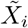 does not have autocorrelations that satisfy the cutoff criteria, the algorithm then checked if the series has a pronounced peak or trough at any time point. The time series’ maximum, 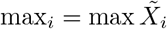, and minimum, 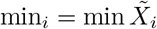 were compared to {max_*cn*_, min_*cn*_}, which are maxima and minima cutoffs from null distributions again computed using the bootstrap time series for all possible time series lengths *n*. 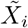 is then labeled as a *SpikeMax* signal if max_*j*_ > max_*cn*_, or as a *SpikeMin* if min_*j*_ < min_*cn*_. A time series that did not meet any of the cutoff criteria was not labeled as having a statistically significant trend.

#### Clustering and heatmaps

After classification, and using PyIOmica’s `clusterTimeSeriesCategorization` function, we carried out a two-tier hierarchical clustering (agglomerative; average linkage) for each temporal class, to identify groups (G) and sub-groups (S). The clustering grouping used a similarity based on *ρ_isn_* (for the autocorrelation classification) and {*Q_is_*} for the second tier. Groups and subgroups were determined using a silhouette algorithm [45]. The results were visualized for each subject and every temporal class identified using PyIOmica’s `visualizeTimeSeriesCategorization`. Example outputs are shown in Fig. 2 and included in the ODFs.

#### Reactome enrichment analysis

Reactome [18] pathway enrichment analysis was carried out for each Group/Subgroup and each subject using the Reactome application programming interface (API) in PyIOmica. Complete output is included in the ODFs.

### Across subject comparisons

#### Network construction

The network analysis was carried out in Python, using `networkx` [46] and `scikit-network` [47]. First the time series periodograms for all the omics time series were computed for all subjects using the `LombScargle` function in PyIOmica. Next, for pairs of subjects *p*, *q* and for each omics time series *i*, a Euclidean distance matrix *D_i_* was constructed. In parallel, a bootstrap simulation of 50,000 time series was also generated from the data, as a null distribution, and the pairwise Euclidean distances were computed for these as well to determine a distance cutoff for significance, *d_c_* at the 0.99 quantile level. Entries were kept that were most proximal to each other, by creating a restricted distance matrix *R_i_*, such that

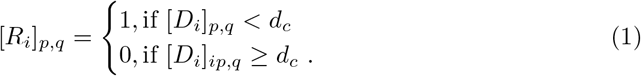

A weighted network was then constructed, with the subjects represented as nodes, with an adjacency matrix *A*, constructed as *A* = ∑_*i*_*R_i_*. The entries *p*, *q* of the adjacency matrix represent the connections in the network. A nonzero entry *A_p,q_* means there is an edge connecting nodes (subject) p and q. The magnitude of *A_p,q_* provides a weight of the edge. In summary, the edges connecting pairs of nodes (subjects) were added to the network if there was at least one omics for which the distance between the subjects was within the cutoff *d_c_*, and the edges were weighted by the number of omics that met this criterion for each pair of subjects.

#### Network communities calculation

To determine the network community structure, a k-means approach was used. The computation used scikit-network’s `clustering.kmeans`, applying an embedding, and utilizing singular value decomposition with dimension one. The number of communities (3) was selected based on the silhouette method [45], and the `sklearn.metrics.silhouette_score` [48] was used for the silhouette scores calculation.

#### Mann-Whitney U tests

Subject measurements were compared between members of the communities in the network calculations (see above). We used non-parametric Mann-Whitney U tests [23], to test for statistically significant pairwise differences across communities (p-value < 0.05). The results shown in Table 3 were computed using the `scipy.stats.mannwhitneyu` Python functionality [49].

## Data and code availability

The original data analyzed in this investigation were made publicly available by Zhou et al. [14] and Schussler-Fiorenza Rose et al. [4] as described therein, on https://med.stanford.edu/ipop.html. All data files used in this investigation, including original code and results files have been released on Github (https://github.com/gmiaslab/TemporalMultiomicsDiabetes), and also deposited on Zenodo (https://doi.org/10.5281/zenodo.5767983), and are referred to as Online Data Files (ODFs) in the manuscript.

## Acknowledgments

This work was supported by the Translational Research Institute for Space Health through NASA Cooperative Agreement NNX16AO69A (project T0412). CP acknowledges support by NIH R01GM122085.

## References

1. Gambhir SS, Ge TJ, Vermesh O, Spitler R. Toward achieving precision health. Science Translational Medicine. 2018;10(430). doi:10.1126/scitranslmed.aao3612.

2. Snyder M, Zhou W. Big data and health. Lancet Digit Health. 2019;1(6):e252–e254. doi:10.1016/S2589-7500(19)30109-8.

3. Kellogg RA, Dunn J, Snyder MP. Personal Omics for Precision Health. Circ Res. 2018;122(9):1169–1171. doi:10.1161/CIRCRESAHA.117.310909.

4. Schussler-Fiorenza Rose SM, Contrepois K, Moneghetti KJ, Zhou W, Mishra T, Mataraso S, et al. A longitudinal big data approach for precision health. Nat Med. 2019;25(5):792–804. doi:10.1038/s41591-019-0414-6.

5. Hasin Y, Seldin M, Lusis A. Multi-omics approaches to disease. Genome Biol. 2017;18(1):83. doi:10.1186/s13059-017-1215-1.

6. Chen R, Mias GI, Li-Pook-Than J, Jiang L, Lam HY, Chen R, et al. Personal omics profiling reveals dynamic molecular and medical phenotypes. Cell. 2012;148(6):1293–1307. doi:10.1016/j.cell.2012.02.009.

7. Ahadi S, Zhou W, Schüssler-Fiorenza Rose SM, Sailani MR, Contrepois K, Avina M, et al. Personal aging markers and ageotypes revealed by deep longitudinal profiling. Nature Medicine. 2020;26(1):83–90. doi:10.1038/s41591-019-0719-5.

8. Tebani A, Gummesson A, Zhong W, Koistinen IS, Lakshmikanth T, Olsson LM, et al. Integration of molecular profiles in a longitudinal wellness profiling cohort. Nat Commun. 2020;11(1):4487. doi:10.1038/s41467-020-18148-7.

9. Sailani MR, Metwally AA, Zhou W, Rose SMS, Ahadi S, Contrepois K, et al. Deep longitudinal multiomics profiling reveals two biological seasonal patterns in California. Nat Commun. 2020;11(1):4933. doi:10.1038/s41467-020-18758-1.

10. Li X, Dunn J, Salins D, Zhou G, Zhou W, Schussler-Fiorenza Rose SM, et al. Digital Health: Tracking Physiomes and Activity Using Wearable Biosensors Reveals Useful Health-Related Information. PLoS Biol. 2017;15(1):e2001402. doi:10.1371/journal.pbio.2001402.

11. Tasaki S, Suzuki K, Kassai Y, Takeshita M, Murota A, Kondo Y, et al. Multi-omics monitoring of drug response in rheumatoid arthritis in pursuit of molecular remission. Nature communications. 2018;9(1):1–12. doi:10.1038/s41467-018-05044-4.

12. Mias GI, Singh VV, Rogers LRK, Xue S, Zheng M, Domanskyi S, et al. Longitudinal saliva omics responses to immune perturbation: a case study. Sci Rep. 2021;11(1):710. doi:10.1038/s41598-020-80605-6.

13. Price ND, Magis AT, Earls JC, Glusman G, Levy R, Lausted C, et al. A wellness study of 108 individuals using personal, dense, dynamic data clouds. Nat Biotechnol. 2017;35(8):747–756. doi:10.1038/nbt.3870.

14. Zhou W, Sailani MR, Contrepois K, Zhou Y, Ahadi S, Leopold SR, et al. Longitudinal multi-omics of host-microbe dynamics in prediabetes. Nature. 2019;569(7758):663–671. doi:10.1038/s41586-019-1236-x.

15. Domanskyi S, Piermarocchi C, Mias GI. PyIOmica: longitudinal omics analysis and trend identification. Bioinformatics. 2019;36(7):2306–2307. doi:10.1093/bioinformatics/btz896.

16. Mias GI, Yusufaly T, Roushangar R, Brooks LR, Singh VV, Christou C. MathIOmica: An Integrative Platform for Dynamic Omics. Sci Rep. 2016;6:37237. doi:10.1038/srep37237.

17. Zheng M, Domanskyi S, Piermarocchi C, Mias GI. Visibility graph based temporal community detection with applications in biological time series. Scientific Reports. 2021;11(1):1–12. doi:10.1038/s41598-021-84838-x.

18. Croft D, O’Kelly G, Wu G, Haw R, Gillespie M, Matthews L, et al. Reactome: a database of reactions, pathways and biological processes. Nucleic Acids Research. 2010;39(suppl_1):D691–D697. doi:10.1093/nar/gkq1018.

19. Randeria SN, Thomson GJ, Nell TA, Roberts T, Pretorius E. Inflammatory cytokines in type 2 diabetes mellitus as facilitators of hypercoagulation and abnormal clot formation. Cardiovascular diabetology. 2019;18(1):1–15. doi:10.1186/s12933-019-0870-9.

20. Dovio A, Angeli A. Cytokines and type 2 diabetes mellitus. JaMA. 2001;286(18):2233–2233. doi:10.1001/jama.286.18.2233.

21. Miranda TS, Heluy SL, Cruz DF, da Silva HDP, Feres M, Figueiredo LC, et al. The ratios of pro-inflammatory to anti-inflammatory cytokines in the serum of chronic periodontitis patients with and without type 2 diabetes and/or smoking habit. Clinical oral investigations. 2019;23(2):641–650. doi:10.1007/s00784-018-2471-5.

22. Fay MP, Proschan MA. Wilcoxon-Mann-Whitney or t-test? On assumptions for hypothesis tests and multiple interpretations of decision rules. Statistics Surveys. 2010;4(none):1 – 39. doi:10.1214/09-SS051.

23. Mann HB, Whitney DR. On a Test of Whether one of Two Random Variables is Stochastically Larger than the Other. The Annals of Mathematical Statistics. 1947;18(1):50 – 60. doi:10.1214/aoms/1177730491.

24. Pan X, Kaminga AC, Wen SW, Liu A. Chemokines in Prediabetes and Type 2 Diabetes: A Meta-Analysis. Frontiers in Immunology. 2021;12:934. doi:10.3389/fimmu.2021.622438.

25. Herder C, Baumert J, Thorand B, Koenig W, De Jager W, Meisinger C, et al. Chemokines as risk factors for type 2 diabetes: results from the MONICA/KORA Augsburg study, 1984-2002. Diabetologia. 2006;49(5):921–929. doi:10.1007/s00125-006-0190-y.

26. Stanya KJ, Jacobi D, Liu S, Bhargava P, Dai L, Gangl MR, et al. Direct control of hepatic glucose production by interleukin-13 in mice. The Journal of clinical investigation. 2012;123(1). doi:10.1172/JCI64941.

27. Vargas-Alarcon G, Perez-Hernandez N, Rodriguez-Perez JM, Fragoso JM, Posadas-Romero C, Lopez-Bautista F, et al. Interleukin 27 polymorphisms, their association with insulin resistance and their contribution to subclinical atherosclerosis. The GEA Mexican study. Cytokine. 2019;114:32–37. doi:10.1016/j.cyto.2018.11.028.

28. Wang R, Han G, Wang J, Chen G, Xu R, Wang L, et al. The pathogenic role of interleukin-27 in autoimmune diabetes. Cellular and molecular life sciences. 2008;65(23):3851–3860. doi:10.1007/s00018-008-8540-1.

29. Wang Q, Li D, Cao G, Shi Q, Zhu J, Zhang M, et al. IL-27 signalling promotes adipocyte thermogenesis and energy expenditure. Nature. 2021;doi:10.1038/s41586-021-04127-5.

30. Shen J, Fang Y, Zhu H, Ge W. Plasma interleukin-22 levels are associated with prediabetes and type 2 diabetes in the Han Chinese population. Journal of diabetes investigation. 2018;9(1):33–38. doi:10.1111/jdi.12640.

31. Shen S, Wang F, Fernandez A, Hu W. Role of platelet-derived growth factor in type II diabetes mellitus and its complications. Diabetes and Vascular Disease Research. 2020;17(4):1479164120942119. doi:10.1177/1479164120942119.

32. Yeboah J, Sane DC, Crouse JR, Herrington DM, Bowden DW. Low plasma levels of FGF-2 and PDGF-BB are associated with cardiovascular events in type II diabetes mellitus (diabetes heart study). Disease markers. 2007;23(3):173–178. doi:10.1155/2007/962892.

33. Heuer JG, Harlan SM, Yang DD, Jaqua DL, Boyles JS, Wilson JM, et al. Role of TGF-alpha in the progression of diabetic kidney disease. American Journal of Physiology-Renal Physiology. 2017;312(6):F951–F962. doi:10.1152/ajprenal.00443.2016.

34. Qiao YC, Chen YL, Pan YH, Ling W, Tian F, Zhang XX, et al. Changes of transforming growth factor beta 1 in patients with type 2 diabetes and diabetic nephropathy: a PRISMA-compliant systematic review and meta-analysis. Medicine. 2017;96(15). doi:10.1097/MD.0000000000006583.

35. Mou X, Zhou DY, Zhou DY, Ma JR, Liu YH, Chen HP, et al. Serum TGF-*β*1 as a biomarker for type 2 diabetic nephropathy: A meta-analysis of randomized controlled trials. PLoS One. 2016;11(2):e0149513. doi:10.1371/journal.pone.0149513.

36. Chen SCC, Tsai SP, Jhao JY, Jiang WK, Tsao CK, Chang LY. Liver fat, hepatic enzymes, alkaline phosphatase and the risk of incident type 2 diabetes: a prospective study of 132,377 adults. Scientific reports. 2017;7(1):1–9. doi:10.1038/s41598-017-04631-7.

37. Malo MS. A high level of intestinal alkaline phosphatase is protective against type 2 diabetes mellitus irrespective of obesity. EBioMedicine. 2015;2(12):2016–2023. doi:10.1016/j.ebiom.2015.11.027.

38. Panee J. Monocyte Chemoattractant Protein 1 (MCP-1) in obesity and diabetes. Cytokine. 2012;60(1):1–12. doi:10.1016/j.cyto.2012.06.018.

39. So BI, Song YS, Fang CH, Park JY, Lee Y, Shin JH, et al. G-CSF prevents progression of diabetic nephropathy in rat. PLoS One. 2013;8(10):e77048. doi:10.1371/journal.pone.0077048.

40. Higurashi M, Ohya Y, Joh K, Muraguchi M, Nishimura M, Terawaki H, et al. Increased urinary levels of CXCL5, CXCL8 and CXCL9 in patients with Type 2 diabetic nephropathy. Journal of diabetes and its complications. 2009;23(3):178–184. doi:10.1016/j.jdiacomp.2007.12.001.

41. Seijkens T, Kusters P, Engel D, Lutgens E. CD40–CD40L: Linking pancreatic, adipose tissue and vascular inflammation in type 2 diabetes and its complications. Diabetes and Vascular Disease Research. 2013;10(2):115–122. doi:10.1177/1479164112455817.

42. Zhang Q, Fang W, Ma L, Wang ZD, Yang YM, Lu YQ. VEGF levels in plasma in relation to metabolic control, inflammation, and microvascular complications in type-2 diabetes: a cohort study. Medicine. 2018;97(15). doi:10.1097/MD.0000000000010415.

43. Feng ZC, Riopel M, Popell A, Wang R. A survival Kit for pancreatic beta cells: Stem cell factor and c-Kit receptor tyrosine kinase. Diabetologia. 2015;58(4):654–665. doi:10.1007/s00125-012-2566-5.

44. Sell H, Eckel J. Chemotactic cytokines, obesity and type 2 diabetes: in vivo and in vitro evidence for a possible causal correlation?: Symposium on ‘Frontiers in adipose tissue biology’. Proceedings of the nutrition society. 2009;68(4):378–384. doi:10.1017/S0029665109990218.

45. Rousseeuw PJ. Silhouettes: A graphical aid to the interpretation and validation of cluster analysis. Journal of Computational and Applied Mathematics. 1987;20:53–65. doi:10.1016/0377-0427(87)90125-7.

46. Hagberg AA, Schult DA, Swart PJ. Exploring Network Structure, Dynamics, and Function using NetworkX. In: Varoquaux G, Vaught T, Millman J, editors. Proceedings of the 7th Python in Science Conference. Pasadena, CA USA; 2008. p. 11 – 15. Available from: http://conference.scipy.org/proceedings/SciPy2008/paper_2/ [cited Nov. 10, 2021].

47. Bonald T, de Lara N, Lutz Q, Charpentier B. Scikit-network: Graph Analysis in Python. vol. 21; 2020. p. 1–6. Available from: http://jmlr.org/papers/v21/20-412.html [cited Nov. 10, 2021].

48. Pedregosa F, Varoquaux G, Gramfort A, Michel V, Thirion B, Grisel O, et al. Scikit-learn: Machine Learning in Python. vol. 12; 2011. p. 2825–2830. Available from: http://jmlr.csail.mit.edu/papers/v12/pedregosa11a.html [cited Nov. 10, 2021].

49. Virtanen P, Gommers R, Oliphant TE, Haberland M, Reddy T, Cournapeau D, et al. SciPy 1.0: Fundamental Algorithms for Scientific Computing in Python. Nature Methods. 2020;17:261–272. doi:10.1038/s41592-019-0686-2.

